# Machine Learning-Assisted Raman Spectroscopy for Quantitative Prediction of Multicomponent Mixtures

**DOI:** 10.64898/2026.11.18.767617

**Authors:** Aman Yadav, Aahil Khambhawala, Duyen Dinh, Hung-Jen Wu

## Abstract

Machine Learning (ML)-assisted Raman spectroscopy that can efficiently analyze molecularly specific fingerprints is well suited for quantitative analysis of complex chemical mixtures. However, developing accurate predictive ML models for multicomponent systems remains difficult because collecting sufficient experimental calibration data is slow, expensive, and limited. In this work, we present a ML-assisted Raman spectroscopy framework for predicting the quantitative composition of multicomponent lipid mixtures (consisting of 15 lipids). To reduce the experimental burden of generating extensive multicomponent spectral datasets, we computationally constructed a large spectral library using experimentally measured pure-component Raman spectra from 15 lipids, including triglycerides, medium- and long-chain fatty acids, and fatty acid methyl and ethyl esters. Mixture compositions were sampled across a constrained compositional space and used to generate training spectra from weighted combinations of the pure component spectra. Experimentally prepared mixtures were then used to evaluate the model under real measurement conditions. To reduce the mismatch between library-generated spectra and experimental mixture spectra, a linear correction strategy was applied to account for systematic spectral differences caused by mixing effects, intermolecular interactions, and measurement variability. The corrected experimental spectra were then used for concentration prediction with an uncertainty-aware workflow. The compositions of the 15 lipid mixtures were successfully predicted using the ML model trained solely on a simulated dataset, achieving *R*^2^ values exceeding 0.95 for each lipid. By combining scalable, library-based model training with experimental validation on physical multicomponent mixtures, this approach minimizes the need for exhaustive calibration datasets while maintaining high quantitative accuracy. The proposed framework offers a practical path for quantitative Raman analysis of chemical mixtures and can easily be extended to quality control applications across the food, pharmaceutical, nutraceutical, and chemical industries.

**Table of Contents Graphic:** 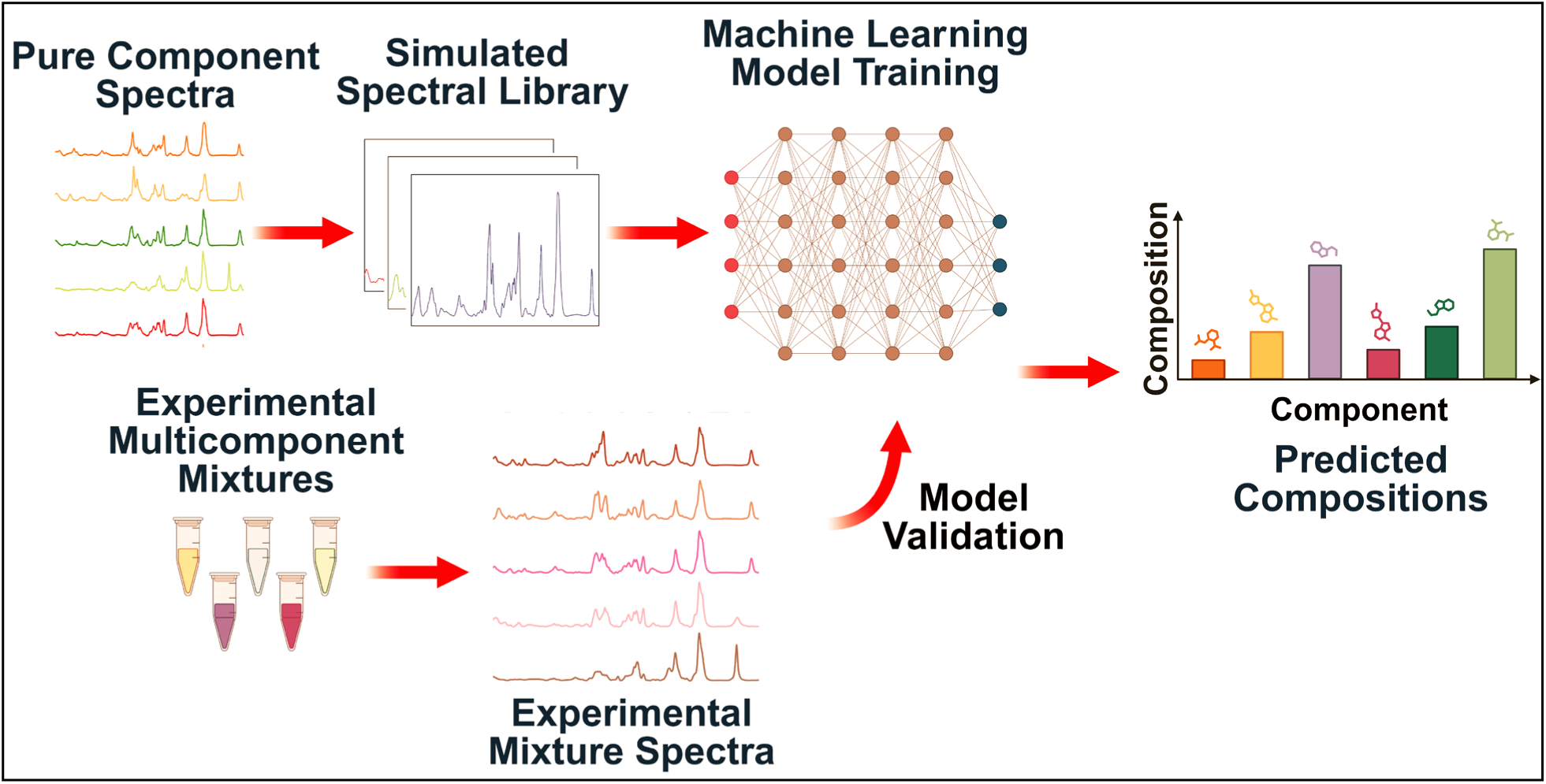

## Introduction

The combination of machine learning (ML) and Raman spectroscopy has emerged as a powerful analytical approach. Raman spectroscopy provides detailed molecular fingerprints, while ML enables efficient processing and interpretation of complex spectral data. This integrated approach has been widely applied across diverse fields for qualitative classification and quantitative analysis, including materials science, biomedical research, forensic science, pharmaceuticals, and food analysis ^1–6^.

Although numerous studies have demonstrated the efficacy of ML-assisted Raman spectroscopy for qualitative classification, the quantitative prediction of analyte composition remains in an early stage of development. For example, Raman spectroscopy combined with chemometric analysis has been used to measure fatty acid composition in adipose tissue and cooking oils^7,8^. Similarly, Tanemura et al. monitored metabolite and culture-component concentrations in cell culture.^9^ Hu et al. used ML-assisted surface-enhanced Raman spectroscopy (SERS) to quantify sialylation levels in glycoproteins.^10^ Additional applications include analyzing mixtures of organic solvents, amino acids, food additives, etc.^11–18^ However, most prior studies have either been limited to relatively small sets of mixtures or lacked external validation using independent experimental datasets. Consequently, a robust protocol capable of analyzing large, complex mixtures has yet to be established.

Quantitative analysis using ML-assisted Raman spectroscopy remains constrained by restricted access to sufficiently large, diverse, and accurately labeled datasets^16^. Generating such datasets experimentally often requires considerable time, cost, and effort.^19^ Consequently, the primary bottleneck in many cases is not the selection of model architecture, but the lack of experimentally prepared mixtures with precisely known compositions for training and validation. To address this challenge, prior studies have demonstrated that the spectrum of a mixture can be approximated as a linear combination of its individual pure-component spectra.^20, 21^ Leveraging this principle, a simulated spectral library can be generated by linearly combining pure-component spectra.

In this work, we propose a strategy to generate a simulated Raman spectral library, apply appropriate spectrum correction methods, and use the corrected data to quantitatively analyze experimentally prepared mixtures (EPMs). This methodology enables the construction of large, diverse, and compositionally well-annotated spectral libraries, thereby reducing the experimental burden of generating extensive multicomponent spectral datasets and supporting the development of more robust and generalizable analytical models. To demonstrate this strategy, we prepared multicomponent mixtures using 15 lipids, including triglycerides and fatty acids (detailed structural representations are provided in Figure S1). These compounds were selected for their high relevance across pharmaceuticals, cosmetics, food products, nutraceuticals, and drug delivery systems.^22, 23^

The approach consists of two main steps. First, pure-component Raman spectra were experimentally acquired (Figure 1). For the simulated spectra, mixture compositions were sampled from a Dirichlet distribution subject to predefined compositional constraints, enabling systematic exploration of the bounded compositional simplex. These compositions were then used to generate synthetic Raman spectra through a linear spectral mixing model, in which each simulated spectrum was constructed as a composition-weighted sum of the pure-component Raman spectra.

**Figure 1.**
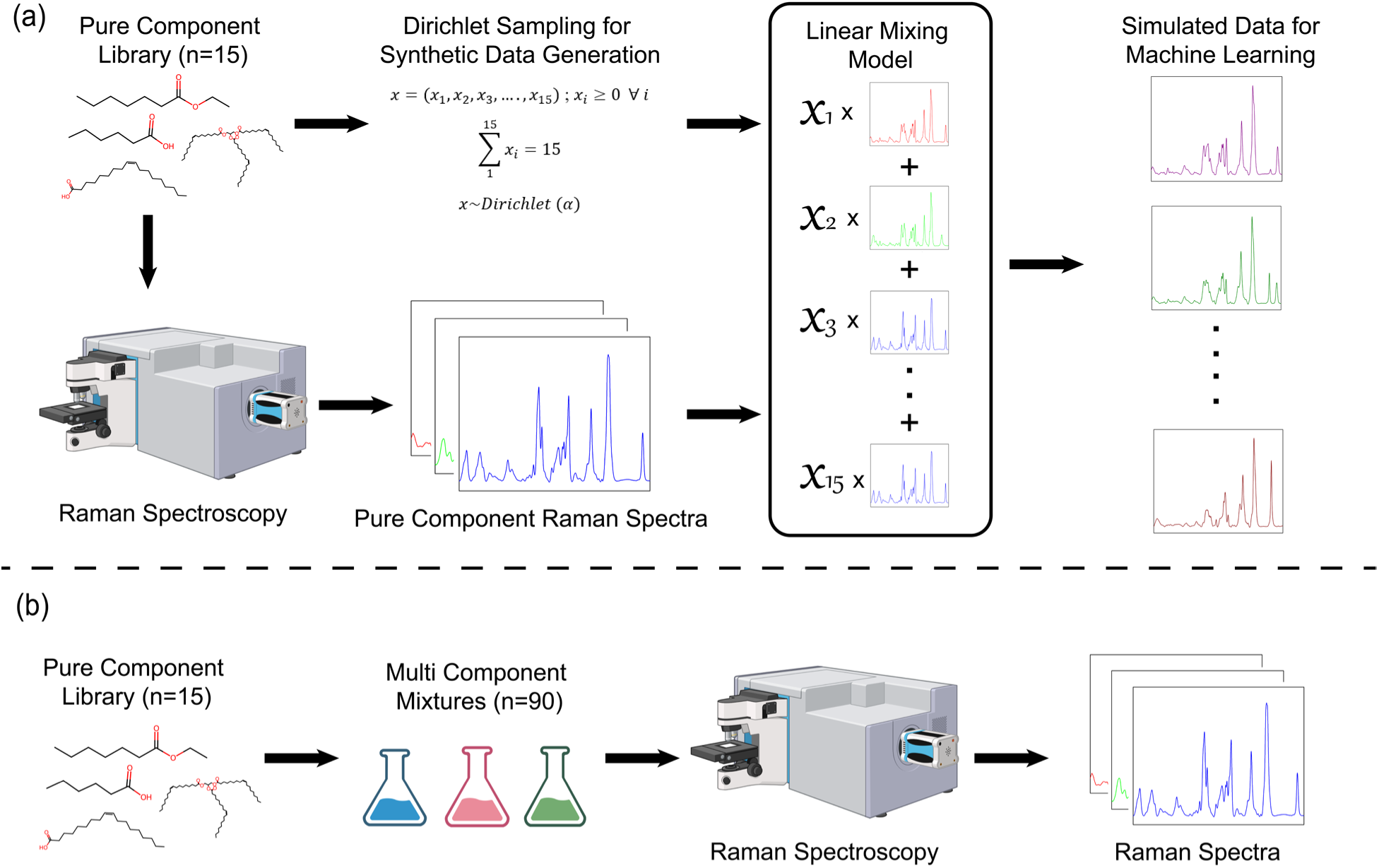
Schematic overview of the dataset-generation workflow. (a) Pure Raman spectra were collected for 15 lipid components. Mixture compositions were then sampled from a constrained Dirichlet distribution, and a simulated spectral library was generated using a linear spectral mixing model. (b) Experimental multicomponent mixtures were prepared across selected compositions, followed by Raman spectral acquisition for experimental validation.

Second, to account for intermolecular interactions that alter molecular vibrations within mixtures, a linear regression (LR)-based correction model was applied to align experimental spectra with simulated data (Figure 2a). Correcting the experimental spectra before prediction helps reduce the domain difference between simulated training data and real experimental measurements. This allows the model to use the large, simulated training set while still accounting for spectral changes caused by real mixture environments.^24–26^ Moreover, a heteroscedastic Gaussian regression model is implemented to estimate the prediction uncertainty of lipid composition (Figure 2b).

**Figure 2.**
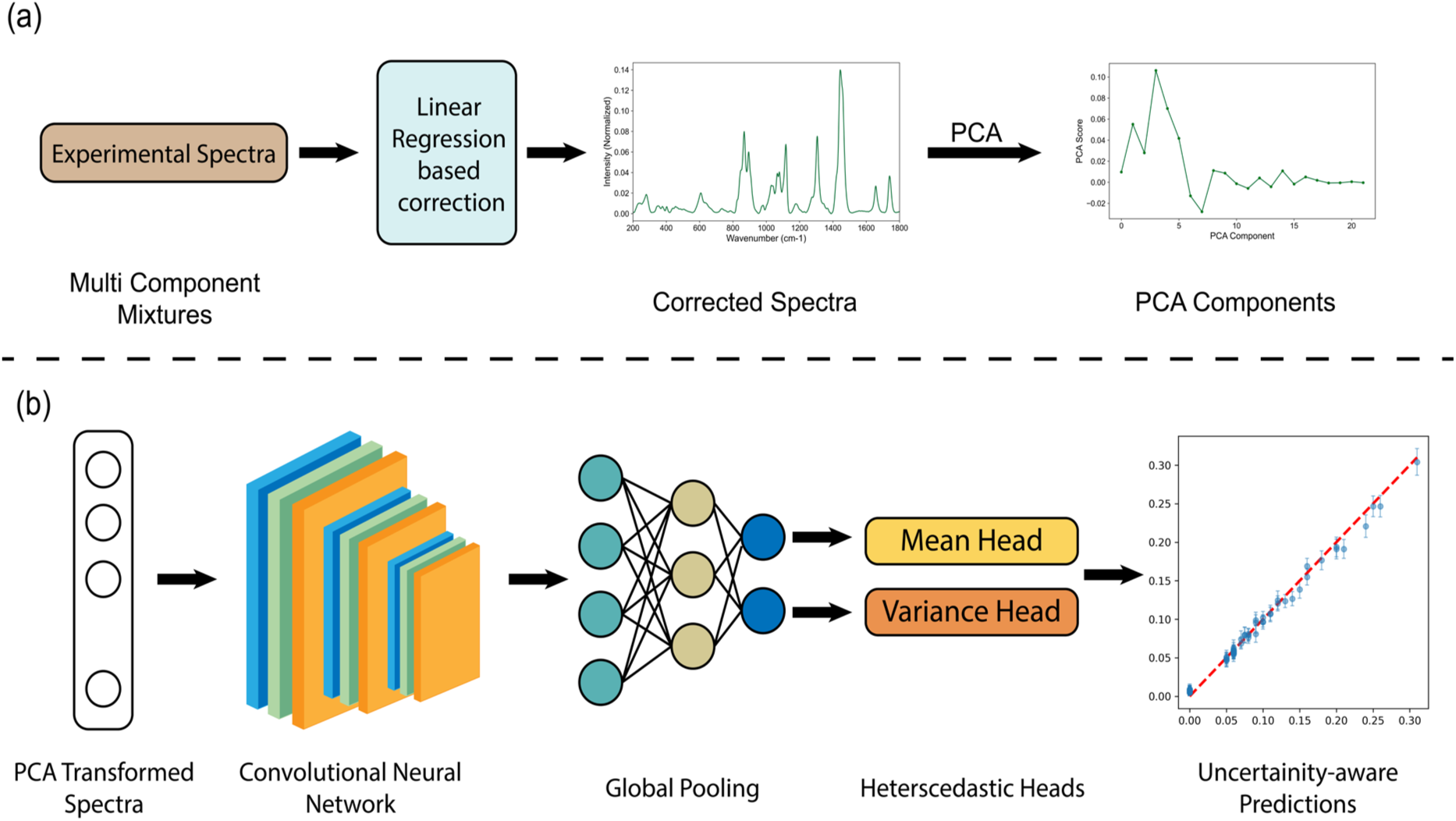
Schematic workflow for data preprocessing and prediction. (a) Simulated and experimental Raman spectra are preprocessed, followed by linear correction for the experimental spectra only and principal component analysis (PCA). (b) PCA-transformed data are then used for uncertainty-aware predictions.

Raman spectra of the 15-lipid EPMs were experimentally acquired for validation. Using a machine learning model trained exclusively on simulated data, our protocol accurately predicted the composition of 15 lipid mixtures (*R*^2^ > 0.95 per lipid). This framework offers a practical pathway for quantitative Raman analysis and broader chemometric applications.

## Methods

### Materials

Glyceryl trioctanoate (≥97%) (GTO), glyceryl trioleate (≥97%) (GTL), hexanoic acid (≥99%) (HxA), heptanoic acid (≥99%) (HpA), methyl hexanoate (99%) (MHx), methyl heptanoate (99%) (MHp), and ethyl heptanoate (99%) (EHp) were purchased from Sigma-Aldrich. Glyceryl tricaproate (>95%) (GTC), glyceryl tributyrate (>97%) (GTB), and oleic acid (99%) (OlA) (manufactured by Tokyo Chemical Industry (TCI)) were purchased from Thermo Fisher Scientific. Octanoic acid (99%) (OcA), methyl octanoate (99%) (MOc), ethyl octanoate (99%) (EOc), ethyl hexanoate (99%) (EHx), and linoleic acid (99%) (LnA) were purchased from Thermo Fisher Scientific. All reagents were used as received without further purification. Raman measurements of fatty acids were conducted using Corning™ 96-well half-area high-content imaging glass-bottom microplates.

### Selected lipids

The selected 15 fatty acid derivatives can be categorized into three groups: triglycerides (glyceryl trioctanoate (GTO), glyceryl trioleate (GTL), glyceryl tricaproate (GTC), and glyceryl tributyrate (GTB)); medium-chain fatty acids and their corresponding methyl and ethyl esters (hexanoic acid (HxA), heptanoic acid (HpA), octanoic acid (OcA), methyl hexanoate (MHx), methyl heptanoate (MHp), methyl octanoate (MOc), ethyl hexanoate (EHx), ethyl heptanoate (EHp), and ethyl octanoate (EOc)); and long-chain fatty acids (linoleic acid (LnA) and oleic acid (OlA)). (structural representations are provided in Figure S1) Lipids such as triglycerides, free fatty acids, and fatty acid esters are important classes of molecules in the nutraceutical, pharmaceutical, and biotechnology industries.^23^ Medium-chain triglycerides, including glyceryl trioctanoate (GTO) and glyceryl tricaproate (GTC), are widely studied for their therapeutic purposes, and due to their shorter chain length, which enables rapid digestion, absorption, and metabolism, making them valuable ingredients for the pharmaceutical and food industry.^27^ Glyceryl tributyrate has shown stable butyrate release, which exerts antiproliferative actions on colon cancer cells.^28^ Glyceryl trioleate (GTL) is also a useful lipid model because oleate-containing triglycerides are abundant in edible oils and provide representative unsaturated lipid structures for studying oxidation, hydrolysis, and lipid-surface interactions. Medium-chain fatty acids and their methyl and ethyl esters, including hexanoic acid (HxA), heptanoic acid (HpA), octanoic acid (OcA), methyl hexanoate (MHx), methyl heptanoate (MHp), methyl octanoate (Moc), ethyl hexanoate (EHx), ethyl heptanoate (EHp), and ethyl octanoate (EOc), are important in flavor, fragrance, fermentation, and fuel-related applications, due to their chain length and esterification affecting volatility, hydrophobicity, and aroma activity.^29^ Ethyl esters are essential aromatic compounds that are generated during microbial fermentation and contribute to the flavor profiles of beverages such as beer and wine.^30^ Long-chain fatty acids such as linoleic acid (LnA) and oleic acid (OlA) are biologically significant. LnA is the most abundant polyunsaturated fatty acid in the human diet, and studies have shown that neurological and skin defects can be reversed by LnA supplementation.^31^ OlA is an important monounsaturated fatty acid. OlA-rich oils, such as olive oil, have been associated with improved plasma lipid and lipoprotein profiles, reduced inflammation, oxidative stress, and coagulation, as well as better glucose homeostasis and blood pressure regulation.^32^

### Sample Preparation

Multicomponent mixtures containing 15 fatty acids, each with different compositions, were prepared as 1 mL solutions. To ensure homogeneity, the samples were thoroughly vortex mixed. We added 150 µL of the prepared samples to the microplate wells for Raman analysis.

### Raman Measurements

A Thermo Fisher Scientific DXR Raman microscope was used to collect Raman spectra with a 780 nm laser excitation at 20 mW. The instrument is equipped with an Olympus BX41 optical microscope and a thermoelectrically cooled charge-coupled detector (Thermo Fisher front-illuminated CCD system) with a 1024 × 256-pixel format, operating at −70 °C. The signal was calibrated using an internal polystyrene standard and a 10× objective (numerical aperture (NA) 0.25). The spot size was approximately 3.8 µm (Airy disk diameter). For pure fatty acids, 20 spectra were collected with an exposure time of 20 seconds for two accumulations at different spots, and for each sample, 10 spectra were collected with the same data acquisition time at different spots.

### Simulated Spectral Library Generation

We generated random volumetric compositions with 15 components using Dirichlet sampling, subject to the constraint that no individual component exceeded a maximum volume fraction (0.5) and each component was either absent (0) or present at a minimum volume fraction of 0.05; intermediate values between 0 and 0.05 were excluded. The constraints were derived from the experimental dataset, and the low-volume fractions (0 to 0.05) were excluded from the experimental mixture design to minimize pipetting error. This constraint ensures that the simulated compositions remain within the experimentally accessible region of compositional space. To generate diverse mixtures, compositions were sampled from a Dirichlet distribution, which naturally produces nonnegative vectors that sum to unity.

We used an approach similar to that discussed in previously published work.^2^ Briefly, we created spectra of fatty acid mixtures by weighted mixing of raw spectra of pure fatty acids.^33, 34^

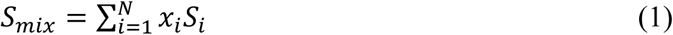

where *S_mix_* denotes simulated spectrum generated using linear mixing, Si denotes its pure-component Raman spectrum, *N* is the total number of components and *x_i_* is the mixing ratio (volume fraction of each component in the mixture).

To comprehensively sample a broad range of compositions, we employed Dirichlet sampling to generate mixing ratios.

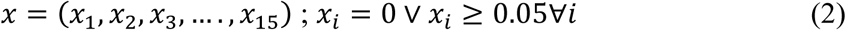

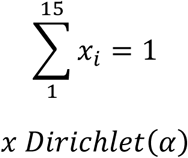

where α is the vector of concentration parameters.

For each simulated mixture, the component fractions were sampled from a symmetric Dirichlet distribution. The same Dirichlet concentration parameter was applied to all 15 molecular components so that no component was favored during mixture generation. The sampled fractions were nonnegative and summed to one, making the Dirichlet distribution appropriate for generating mixture compositions. The concentration parameter controls the type of mixtures generated, where smaller values produce mixtures dominated by fewer components and larger values produce more uniform mixtures. To generate mixtures with different levels of sparsity, the concentration parameter (α) was sampled from a uniform range between 0.5 and 2.0.^35, 36^ Samples that satisfied the maximum-component constraint were accepted directly. If a sampled composition violated the constraint, its components were clipped to the allowable maximum and renormalized and retained only if the constraint was satisfied after renormalization. This approach combines Dirichlet sampling with constraint enforcement to efficiently generate compositions within a bounded simplex. The resulting simulated dataset consists of 100,000 spectra, each represented on a fixed, uniform grid, paired with their corresponding composition vectors, and serves as the primary training data for the machine learning model.

### Spectra Pre-processing

Raw spectra, both simulated and experimental, were interpolated onto a common uniform grid spanning the full spectral range, then cropped to the region of interest (200-1800 cm^-1^). A preprocessing workflow consistent with established methods in the literature was employed.^4, 37^ Baseline correction was applied using an adaptive iteratively reweighted penalized least squares method to remove broad background variations. The spectra were then normalized using vector normalization and shifted to ensure non-negativity, reducing variability due to intensity scaling and baseline offsets. This standardization is critical to prevent the model from learning artifacts related to preprocessing inconsistencies rather than meaningful compositional features.

### Linear Regression (LR) Based Correction

To reduce systematic differences between simulated and experimental spectra, an affine linear correction model was trained using paired spectra from the calibration set. Each calibration sample had an experimentally measured spectrum and a corresponding simulated spectrum generated from the known component composition. The experimental spectra were arranged into a matrix, where each row represented one measured spectrum after preprocessing and interpolation onto the same Raman shift grid. The corresponding simulated spectra were arranged into a second matrix with the same dimensions. The correction model was trained to find a coefficient matrix and intercept vector that transformed the experimental spectra into the simulated spectral domain. The parameters were obtained by minimizing the squared difference between the corrected experimental spectra and the paired simulated spectra. Here, the calibration set contained 50 paired spectra. The trained correction was then evaluated on the remaining 40 experimental samples that were not used to estimate the correction parameters. For a new experimental spectrum, the corrected spectrum was calculated by applying the learned coefficient matrix and intercept vector. This correction was applied only to experimental spectra. The simulated spectra used to train the composition prediction model were already in the target simulated domain and were therefore not adjusted. After correction, the experimental spectra were represented in a spectral space more consistent with the simulated training spectra. The corrected experimental spectra were then used as inputs to the composition prediction model to estimate the fraction of each component in the experimental mixtures.

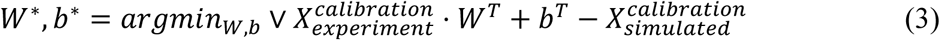

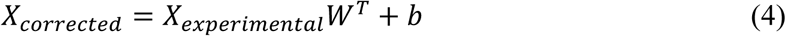

Here W is the weight matrix, b is the bias matrix, and X is the spectra matrix.

### Principal Component Analysis (PCA) Processing

To further standardize the spectral representation and reduce dimensionality, principal component analysis was applied to the spectra prior to model training. The PCA transformation was learned using simulated spectra, ensuring that the feature space reflects the structure of the simulated domain on which the model is trained. The number of retained components was selected to preserve 99.99 percent of the variance in the data, which resulted in a reduced representation of 22 principal components. For simulated training data, spectra were directly projected into the PCA space. For experimental data, spectra were first mapped to the simulated spectral domain using a linear spectral standardization procedure. This procedure is conceptually similar to direct standardization methods used in calibration transfer, where paired spectra are used to learn a linear transformation between two spectral domains. After standardization, the corrected experimental spectra were projected onto the PCA space fitted using the simulated training spectra. This ensures that both simulated and experimental spectra are represented in a consistent feature space before being passed to the neural network. This transformation serves two purposes. First, it reduces the dimensionality of the input spectra, improving computational efficiency and reducing the risk of overfitting. Second, PCA transforms correlated spectral variables into a smaller set of orthogonal principal components ordered by explained variance. In chemometric analysis, PCA has been described as a method for separating systematic data structure from residual noise, where the leading components retain the most useful information and later components mainly contain noise. PCA has also been used in Raman and SERS analysis to preserve the most valuable spectral variance while reducing redundant spectral information.^38–40^

Therefore, retaining the leading principal components provides a compact spectral representation for downstream composition prediction. As a result, the neural network operates on a compact and standardized representation that is aligned across both training and evaluation data.

### Neural Network Model Architecture

A one-dimensional convolutional neural network was used to model the mapping from spectral features to mixture composition. The model operates on the reduced-dimensional representation obtained after principal component analysis, where each input is a 22-dimensional feature vector. The architecture consists of four convolutional layers with channel dimensions 64, 128, 256, and 512, respectively. Each layer applies one-dimensional convolutions with stride-based down-sampling, followed by batch normalization and nonlinear activation. The final feature map is aggregated using global average pooling to obtain a fixed-length representation, which is passed to fully connected output layers. The composition output is obtained using a softmax transformation to enforce nonnegativity and a unit-sum constraint across the 15 components. To estimate prediction uncertainty, the composition model was formulated as a heteroscedastic Gaussian regression model.

During training, the model was optimized using a Gaussian negative log-likelihood loss.

Loss function is given by:

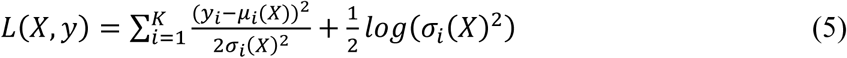

where *x* represents the input Raman spectrum, *y_i_* is the experimentally measured fraction of the *i*^th^component, and *K* is the total number of components in the mixture. The term *μ_i_*(*X*) denotes the predicted mean composition of the *i*^th^component, while 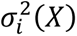 represents the predicted variance, which quantifies the model uncertainty associated with that prediction. µ(*X*) = *softmax* (*f*_µ_(*X*)) and *σ*(*X*) = *softplus*(*f_σ_*(*X*)) where, functions *f_μ_*(*X*)and *f_σ_*(*X*) are the ML model outputs used to estimate the mean composition and uncertainty, respectively.

This loss penalizes large prediction errors while also penalizing unrealistically large or small uncertainty values. Therefore, the model learns both the predicted component concentrations and the reliability of each prediction. During inference, the predicted mean was used as the final concentration estimate, while the predicted standard deviation was used as the uncertainty associated with each component prediction.^41, 42^

### Model Training

The model was trained on simulated spectra generated from composition-weighted combinations of pure component spectra described in simulated spectral library generation. A subset of 25,000 samples was used for model development, with the data randomly partitioned into training and validation sets. Experimental spectra were excluded from training and used only for evaluation to assess generalization to real-world measurements. Training was performed using a heteroscedastic Gaussian regression formulation. In this approach, the model predicts both the mean composition and a positive variance or standard deviation for each component, following mean-variance estimation neural networks proposed by Nix and Weigend.^41^ The model was optimized using equation 5, which is commonly used in this framework to train networks that output both the mean and variance of a normal target distribution. This allows the predicted uncertainty to depend on the input spectrum, so spectra with larger residual mismatch or noise can be assigned larger predicted variance. Because heteroscedastic likelihood training can be sensitive to optimization, the predicted standard deviations were interpreted as model-reported uncertainty estimates rather than guaranteed calibrated confidence intervals ^31, 32^.

In the implementation, the uncertainty head produced one raw scalar value for each of the 15 components. These raw outputs were not used directly as standard deviations because they can be negative. Instead, each raw value was passed through a softplus function and then shifted by a fixed lower bound to obtain the final predicted standard deviation. We used the same value for all components. This lower bound was included only for numerical stability in the Gaussian negative log-likelihood, since division by very small variance values can cause unstable gradients during training (equation 5). The total loss contained the heteroscedastic Gaussian negative log-likelihood and an additional uncertainty penalty. The penalty was computed from the mean predicted standard deviation across the 15 components and weighted by a coefficient (lambda=1e^-4^). This term was included to discourage the model from assigning uniformly large uncertainty values to reduce the contribution of the squared error term. During inference, the same softplus transformation and lower bound were applied to the uncertainty head output, and the resulting standard deviations were reported as component-specific model uncertainty estimates. Optimization was carried out using mini-batch stochastic gradient-based methods, allowing efficient training over the dataset. Model performance was monitored on a held-out validation set throughout training, and the best-performing model was selected based on validation loss. Early stopping was implemented with a patience of 20 epochs to prevent overfitting.

At inference time, experimental spectra were first transformed as mentioned in eqn 4. The trained model was then used to predict composition along with associated uncertainty. Performance was evaluated using standard regression metrics, including mean absolute error (MAE), root mean squared error (RMSE), and coefficient of determination (R^2^), computed across all components.

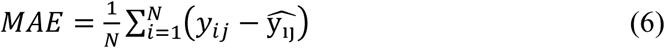

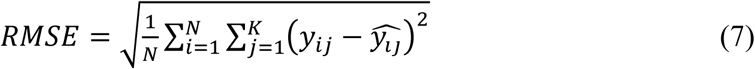

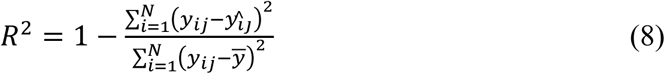

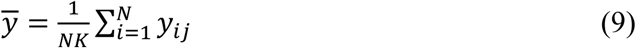

where N is the number of spectra, *y_ij_* is the true fraction of component j in spectrum i, *y*^*_ij_* is the predicted fraction of component j in spectrum i, and *y̅* is the mean true fraction calculated across all spectra and all components.

## Results

### Structurally Similar Lipids Selected for Evaluating the Detection Performance

The selected lipids are both biologically relevant and structurally similar, providing a robust basis for evaluating the limit of detection. (detailed descriptions in Method section) As shown in Figure S2 and S3, the Raman spectra of the selected lipids highlight their strong spectral overlap. The high degree of spectral overlap arises from the shared structural features of the selected lipids, especially long hydrocarbon chains and carbonyl-containing functional groups.^43, 44^ Many Raman bands appear at similar positions across the spectra. For example, the Raman band near 1746 cm^-1^ corresponds to the ester carbonyl stretching, which is present in all triglycerides as well as methyl and ethyl esters. Similarly, the Raman band near 1662 cm^-1^ corresponds to the carboxylic group, which is common to all fatty acids.^45, 46^

The PCA loading plots show that the main sources of spectral variance correspond to chemically meaningful Raman bands associated with lipid functional groups. PC1, which accounts for 45.95% of the total variance, is dominated by a strong positive contribution from the peak at approximately 1660 cm^-^¹ (associated with carboxylic acid groups) and a strong negative contribution near 1746 cm^-^¹ (associated with ester carbonyl vibrations) (Figure S4). This indicates that PC1 primarily distinguishes between fatty acid-rich lipids and ester-containing lipids. PC2, which explains 14.23% of the variance, is driven by features in the fingerprint region (Figure S5). These include bands related to C-C skeletal vibrations, CH₂/CH₃ deformations, hydrocarbon chain conformation, and modes associated with carbonyl groups and unsaturation. Consequently, PC2 captures more subtle structural differences among lipids that share similar hydrocarbon backbones.

PCA loading vectors thus highlight the inherent complexity of the lipid spectral library. The major contributions to both PC1 and PC2 originate from common lipid functional groups, such as hydrocarbon chain vibrations, CH₂/CH₃ deformation modes, unsaturation-related bands, carboxylic acid stretching, and carbonyl stretching. Because these vibrational features are widely shared among triglycerides, fatty acids, and fatty acid esters, there is significant spectral overlap.

To quantify the spectral similarity, we used cosine similarity and the Pearson correlation coefficient (Figure S6 and S7). These metrics have been used in prior studies to quantify similarity in Raman spectra.^47, 48^ Both similarity matrices demonstrate substantial spectral overlap among the selected lipids, with most pairwise similarities exceeding 0.80 and several exceeding 0.90. Particularly high similarities were observed among structurally related compounds, including the ethyl esters EHp, EHx, and EOc (0.96-0.98), the triglycerides GTC and GTO (0.96), and GTL and OlA (0.99). These results indicate that the spectra are dominated by common vibrational features associated with hydrocarbon chains, ester functionalities, and CH_2_/CH_3_ groups. In contrast, LnA and GTB showed comparatively lower similarity to several other components, suggesting more distinct spectral signatures.

Thus, the differentiation of individual lipids does not rely on unique marker peaks but rather on subtle variations in peak intensity, band shape, and relative contributions across these common regions. This extensive spectral overlap underscores the limitations of conventional peak-based or deconvolution methods and justifies the use of multivariate or machine-learning-based approaches for effective resolution.

### Quantitative Prediction of Lipid Composition Using Simulated Raman Spectral Library

Mixture spectra were simulated as linear combinations of pure lipid spectra using constrained Dirichlet-sampled compositions of the 15 lipids. The resulting library was used to train the ML model, which was then validated on a subset of the simulated library. Figure 3 shows the parity plots for comparing the predicted and actual compositions of each lipid for the validation dataset, which is simulated data. The validation results show that the model trained on simulated data has high prediction accuracy with R^2^ values exceeding 0.99 for each lipid.

**Figure 3.**
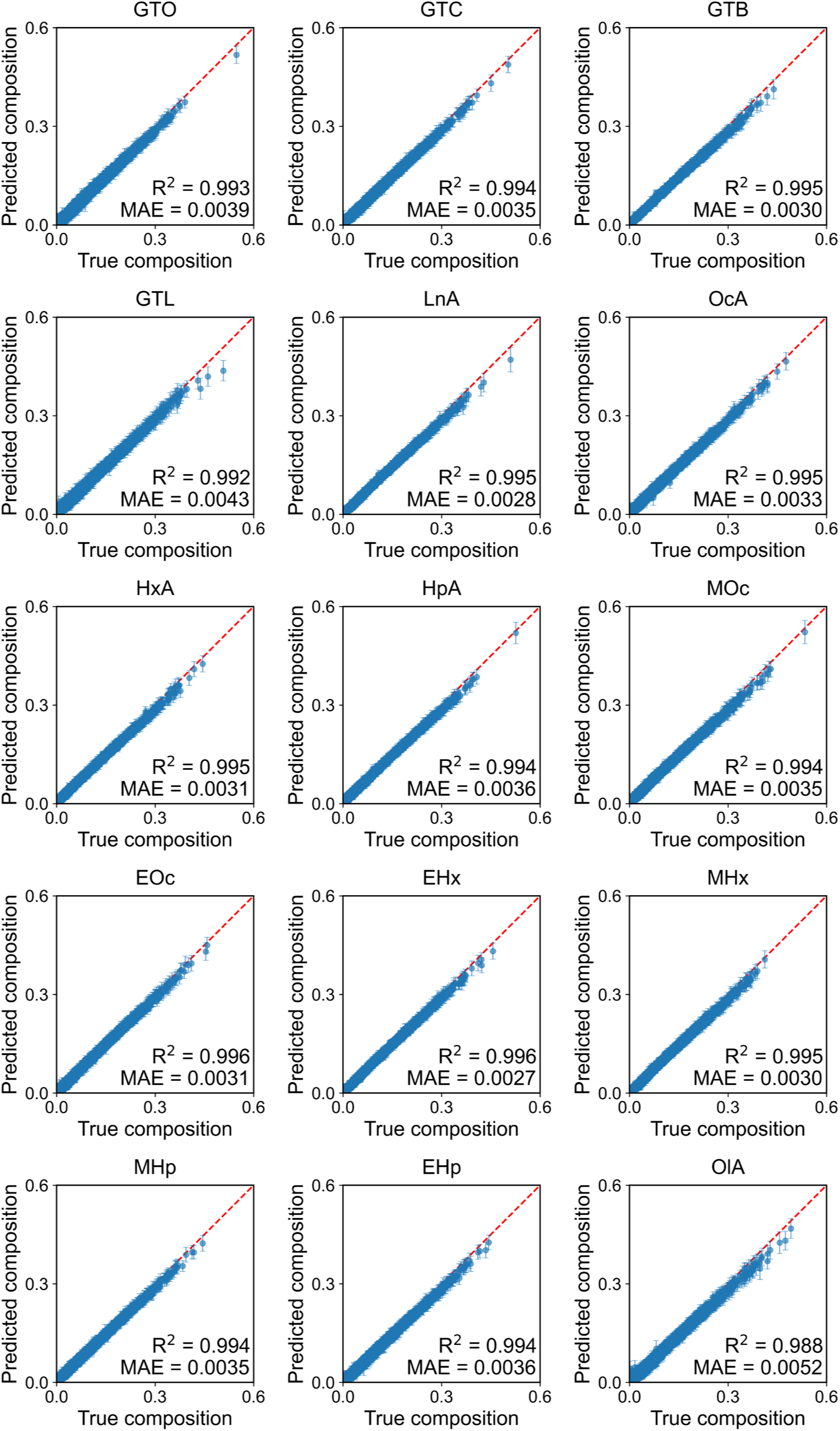
Parity plots for prediction of individual lipid composition of simulated mixtures. The model was trained using the simulated spectral library and validated on simulated dataset. Each point (blue) represents the predicted volume fraction of an individual lipid component in a single EPM, plotted against its actual volume fraction. The dashed diagonal line (red) represents the ideal parity condition, where the predicted and actual compositions are equal. Error bars represent 2 standard deviations of uncertainty associated with the predictions.

To evaluate whether the model trained on simulated data could accurately predict the compositions of experimentally prepared mixtures (EPMs), the model was tested on the EPM dataset. Figure 4 shows parity plots comparing the predicted and actual compositions of each lipid in the EPMs. For most lipids, high R² values were obtained, indicating that the simulated spectral library can be used to predict the compositions of the EPMs. However, a few lipids showed lower prediction accuracy, as reflected by comparatively lower R² values. Among them, GTL, EHp, MOc, and OcA exhibited the lowest R² values. These prediction errors may arise from experimental variability, substantial spectral overlap among structurally similar lipids, and inherent limitations in the model’s ability to resolve closely correlated lipid signals.

**Figure 4.**
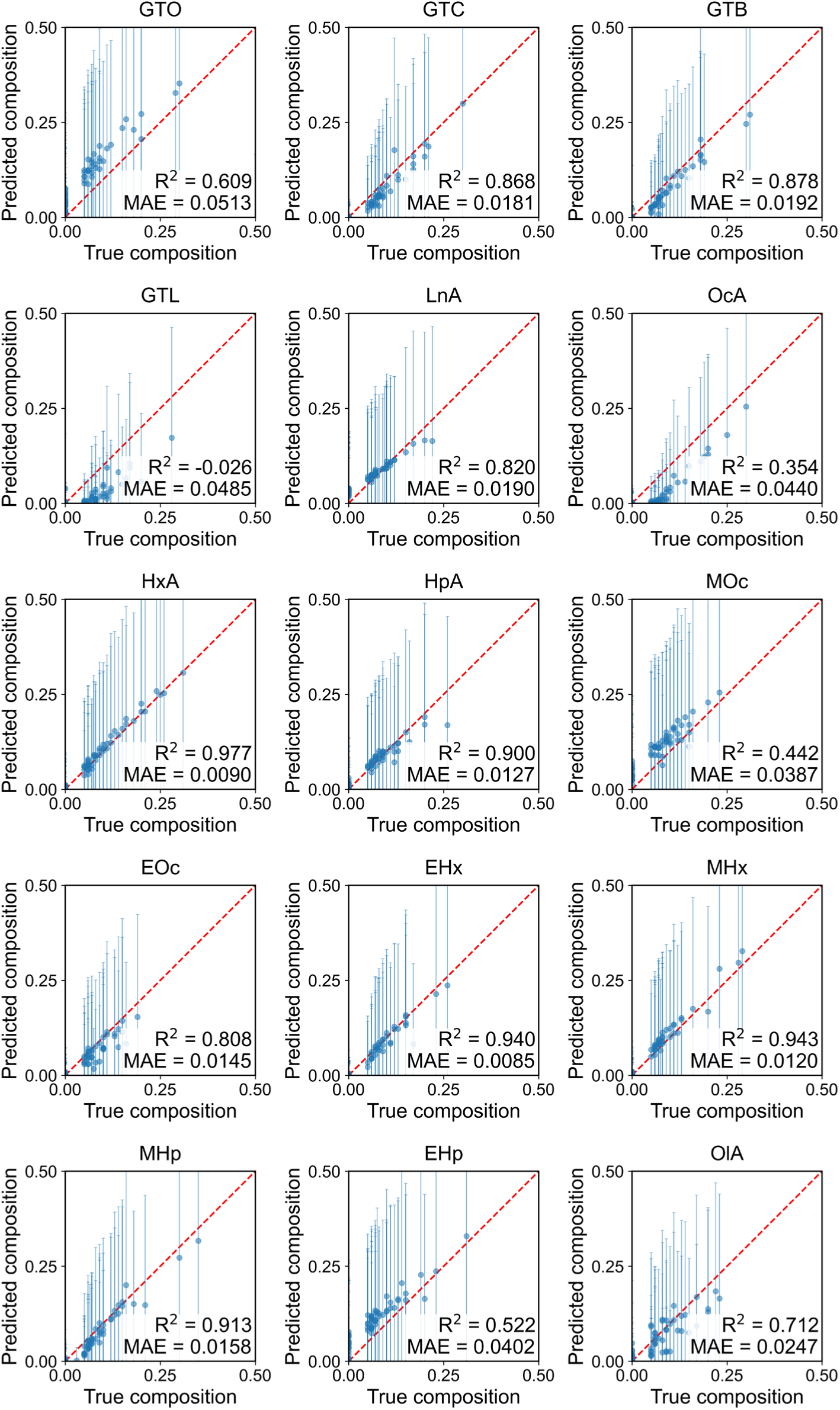
Parity plots for prediction of individual lipid composition in experimentally prepared mixtures (EPMs). The model was trained using the simulated spectral library and validated using Raman spectra of the EPMs. Each point (blue) represents the predicted volume fraction of an individual lipid component in a single EPM, plotted against its actual volume fraction. The dashed diagonal line (red) represents the ideal parity condition, where the predicted and actual compositions are equal. Error bars represent 2 standard deviations of uncertainty associated with the predictions.

Additionally, the predicted uncertainty was further used to construct lipid-wise confidence intervals for the composition estimates^41, 49^. For each lipid, the interval was calculated from the heteroscedastic standard deviation predicted by the model as *μ_i_*(*x*) ± 1.96*σ_i_*(*x*), corresponding to an approximate 95 percent confidence interval under the Gaussian error assumption. These intervals provide an estimate of the reliability of each predicted composition rather than only the point prediction. Narrow intervals indicate regions where the model is confident in the predicted mixture composition, whereas wider intervals indicate greater uncertainty, which may arise from spectral overlap, lower lipid concentration, or differences between simulated and experimental spectra. The agreement between the predicted values and the true compositions, together with the associated confidence intervals, was used to assess both prediction accuracy and uncertainty calibration. The large error bars indicate substantial uncertainty in the predictions, suggesting that the model has low confidence in the predicted compositions.

As discussed previously, based on cosine similarity and Pearson correlation (Figures S6 and S7, respectively), multiple lipids exhibit very similar Raman spectra. Specifically, OlA and GTL have a cosine similarity score and Pearson correlation coefficient of 0.99. We compared the pure-lipid Raman spectra of OlA and GTL (Figure S8). These lipids have closely related molecular structures, as GTL consists of three OlA residues esterified to a glycerol backbone. Consequently, their Raman spectra are highly similar, which may contribute to cross-prediction errors between these lipids.

PCA was further performed to compare simulated and experimental spectra prepared at identical lipid ratios, in order to assess spectral differences between the simulated library and the experimentally measured mixtures (Figure S9). The PCA results, together with spectral comparison, indicate that although linearly mixed simulated spectra provide a controlled and scalable training dataset, they do not fully capture all features of experimentally measured spectra. In practice, experimental spectra exhibit systematic differences relative to simulated spectra due to factors such as instrumental response, measurement noise, baseline variations, intensity distortions, and preprocessing artifacts. Consequently, a model trained solely on simulated spectra would be expected to generalize poorly when evaluated on experimental data, as the two datasets originate from different spectral domains. This mismatch represents a domain shift problem, in which discrepancies in data distribution between the training and evaluation sets can significantly degrade predictive performance.

### Linear Correction Strategy for Experimental Raman Spectra

To address the issue of mismatch, we introduced a spectral domain correction step that maps experimental spectra into the simulated spectral domain prior to model inference. Paired data were constructed by associating each experimental spectrum with a composition-matched simulated spectrum generated from its known composition.

We employed a multi-output linear regression model that was trained to transform experimental spectra into their corresponding simulated spectra. Each experimental spectrum was used as the input, while the matched simulated spectrum was used as the regression target. This allowed the model to learn systematic linear differences between the experimental and simulated spectral domains. The correction model was trained using 50 experimentally measured samples. After training, the linear correction was applied only to the experimental spectra, since the simulated spectra used for model training were already in the simulated domain and did not require further adjustment. Following this correction, the simulated and experimental spectra were represented in a more consistent feature space, improving the alignment between the training and evaluation datasets. Figure 4 compares the spectra and PCA score before and after the linear correction.

To better understand what the linear correction model learned, we visualized the learned weight matrix to examine the spectral regions that contributed most strongly to the correction, as shown in Figure 5. The weight matrix describes the relationship between input and output wavenumbers and reveals how the correction term modifies the experimental spectra to better match the simulated spectra. The strongest positive and negative weights were concentrated in selected Raman regions, indicating that the model learned localized, band-specific corrections. These regions corresponded to the spectral bands showing the largest mismatch between the simulated and experimental spectra, suggesting that the linear correction term captures systematic spectral differences rather than introducing arbitrary changes across the full spectral range (Figure S10).

**Figure 5.**
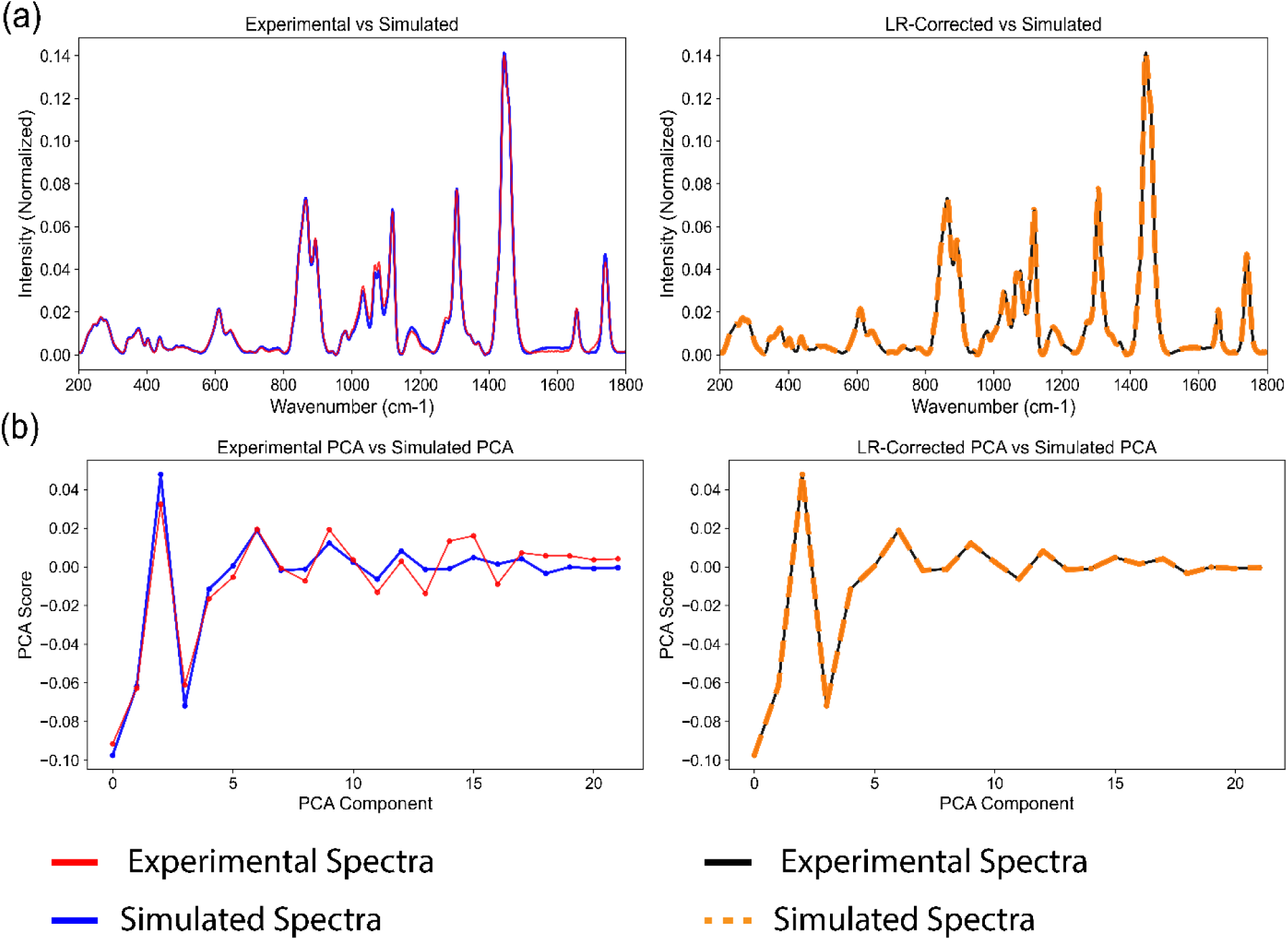
Effect of linear regression (LR)-based correction on the mismatch between experimentally acquired and simulated Raman spectra. (a) Comparison of normalized Raman spectra before and after LR correction. The left panel compares an experimental spectrum with its corresponding simulated spectrum, showing spectral differences. The right panel compares the LR-corrected experimental spectrum with the simulated spectrum, demonstrating substantially improved spectral alignment after correction. (b) Comparison of principal component analysis (PCA) scores of the experimental and simulated spectra. Before correction (left), differences are observed between the PCA score profiles of the experimental and simulated spectra. Following LR correction (right), the corrected experimental PCA scores closely overlap with those of the simulated spectrum across the evaluated principal components, indicating that the correction model effectively transforms the experimental spectra into the simulated spectral domain.

### Quantitative Prediction of Fatty Acid Composition After Spectral Correction

After applying the linear correction to the experimental spectra, the model trained using simulated spectra was used to predict the individual lipid composition of the corrected EPM. Figure 6 shows the parity plot for predicting the composition of each lipid in the EPMs. Following linear correction, the model achieved an R² value greater than 0.95 for each component. Importantly, GTL, EHp, MOc, and OcA, which previously showed the lowest prediction accuracy, exhibited substantial improvement after correction. This indicates that the correction strategy effectively reduced systematic differences between the simulated and experimental spectra, leading to more accurate composition prediction in the EPMs.

**Figure 6.**
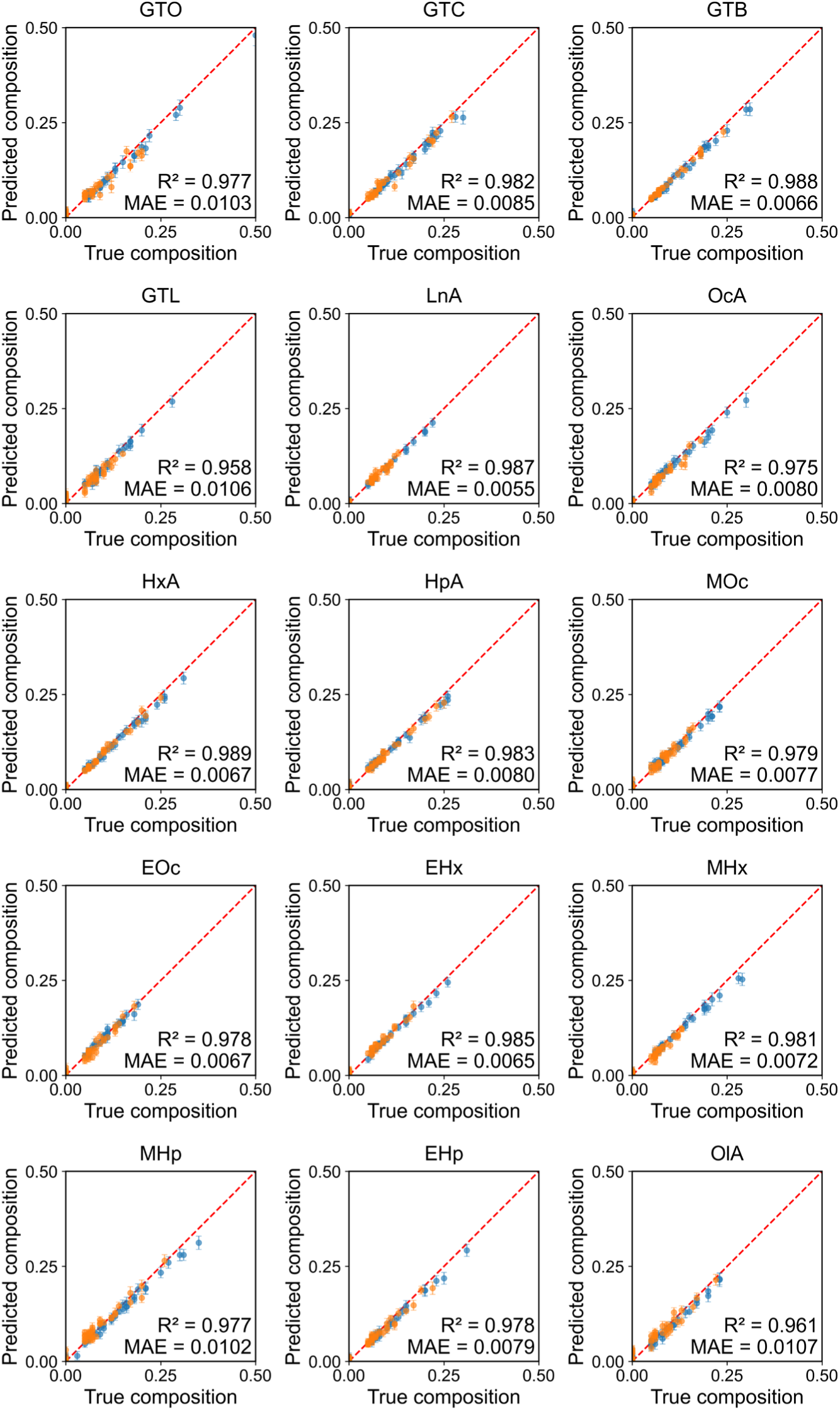
Parity plots showing the simultaneous prediction of individual lipid compositions in experimentally prepared mixtures (EPMs) following LR-based spectral correction. The model was trained using the simulated spectral library and validated using Raman spectra of the EPMs. Each point (orange and blue) represents the predicted volume fraction of an individual lipid component in a single EPM, plotted against its actual volume fraction. Blue points are the points used for training the linear regression-based correction model, while orange points are the points on which the correction model was tested. The dashed diagonal line (red) represents the ideal parity condition, where the predicted and actual compositions are equal. The orange points represent those used to train the LR-correction model. Error bars represent 2 standard deviations of uncertainty associated with the predictions.

The predicted uncertainty was further used to construct component-wise confidence intervals for the composition estimates. The narrower error bars, compared to Figure 4, indicate the model is confident in the predicted mixture composition in the case of the EPMs after correction is applied.

### Influence of Training Dataset Size on Model Performance

The ML model performance is strongly dependent on the size and representativeness of the training dataset. When the training dataset is small, the model may not adequately capture the underlying spectral variability, which can reduce generalization performance and increase the risk of overfitting. To examine the effect of training dataset size on model performance, we trained the model on datasets ranging from 10^3^ to 10^5^ spectra. Figure 7 illustrates how RMSE, MAE, R^2^, and prediction uncertainty vary with the size of the training dataset. As the training size increased from 10^3^ to 10^5^ spectra, RMSE and MAE decreased, while R^2^ increased, indicating improved predictive performance with larger training datasets. The most substantial improvement was observed between 10^4^ and approximately 2×10^4^ spectra, after which the performance began to plateau. The test set showed consistently higher RMSE and MAE and lower R^2^ than the training and validation sets, suggesting reduced generalization to unseen experimental spectra. Overall, these results suggest that increasing the training dataset size improves model accuracy, but the benefit plateaus beyond a certain point.

**Figure 7.**
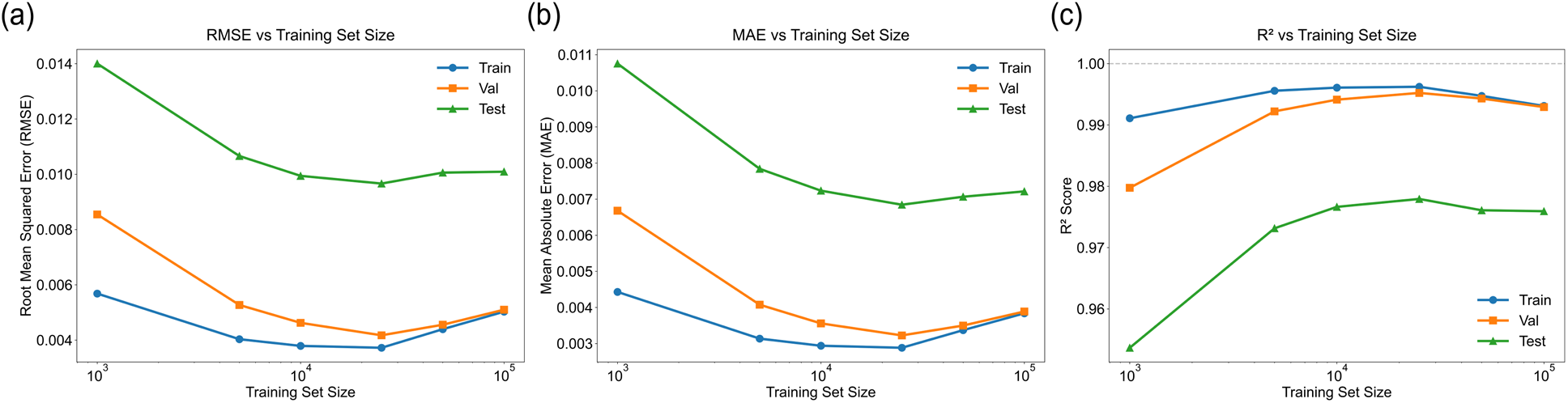
Influence of simulated training dataset size on model performance. Performance was evaluated using (a) root-mean-square error (RMSE), (b) mean absolute error (MAE), and (c) coefficient of determination (R²). Blue, orange, and green curves represent performance on the training, validation, and test datasets, respectively, as a function of training dataset size.

## Discussions

More recent studies have used deep learning models to reduce the need for manual preprocessing and to directly learn the relationship between mixture spectra and component concentrations. CNN-based models have been applied to Raman mixture regression by learning local peak patterns and overlapping spectral features. Studies involving liposome and small-molecule mixtures demonstrated successful regression using synthetic spectra, even with incomplete component libraries. ^10^. Similarly, 1D CNN architectures such as RaMixNet have been developed for mixture classification and regression.^13^ Other approaches include fully connected networks for predicting component contributions in organic solvent mixtures, outperforming DeepRaman and non-negative elastic net on simulated datasets. However, their generalizability to experimental mixtures remains uncertain. Feature fusion networks incorporating attention mechanisms and DCGAN-based data augmentation have also been explored, although some studies lack sufficient experimental details and clearly defined quantitative accuracy metrics.^14^

SERS-based multicomponent studies show a similar trend. Studies have used artificial neural networks to quantify several analytes simultaneously from experimentally measured SERS spectra.^16, 17^ These models are useful because SERS spectra can contain strong substrate effects, nonlinear intensity changes, and peak overlap. However, many SERS studies use small numbers of mixtures or report only relative prediction errors rather than full regression metrics such as R², RMSE, or MAE. Therefore, while these studies demonstrate the potential of ML-assisted Raman and SERS for quantitative mixture analysis, the strength of the validation varies considerably across the literature. The most rigorous quantitative studies are those that include experimentally prepared mixtures with known composition^15^. The study suggests that deep learning is most useful when the spectral response is nonlinear, noisy, or affected by interfering backgrounds, whereas simpler chemometric models may be sufficient for cleaner systems.

Overall, the literature shows a progression from classical chemometric regression toward CNNs, FCNs, attention-based networks, and spectral-extraction neural networks. However, despite these methodological advancements, much of the existing literature lacks rigorous experimental validation. Despite the broad applications of ML-assisted Raman spectroscopy in quantitative and qualitative analysis, data scarcity remains a major challenge because collecting large, diverse, and well-annotated Raman datasets is often difficult, expensive, and time-consuming.^19^ The key limitation in many studies is not the model architecture itself, but the availability of experimentally measured mixtures with known ground truth composition. For this reason, a strong quantitative Raman model should be evaluated using real mixtures, report concentration-level regression metrics, compare against standard chemometric baselines, and assess detection limits for low-abundance components.

In this work, we present a methodology for generating a simulated Raman spectral library, training a predictive model on the simulated spectra, and applying it to the simultaneous multicomponent analysis of experimentally prepared mixtures (EPMs).

Although the simulated spectra closely resembled the Raman spectra of experimentally prepared mixtures with the same compositions, differences can arise between experimental and simulated spectra due to instrumental response, measurement noise, baseline variation, intensity distortion, and preprocessing artifacts. ^2^ Therefore, correcting the experimental spectra before prediction helps reduce the domain difference between simulated training data and real experimental measurements. To reduce this mismatch, we propose a linear regression-based correction model that learns systematic differences between the two datasets. The experimental spectra are corrected based on the simulated spectra. This allows the model to use the large, simulated training set while still accounting for spectral changes caused by real mixture environments.^24–26^

Uncertainty estimation in regression models helps quantify the level of risk inherent in predicted outcomes. A 1-D convolutional neural network model is formulated as a heteroscedastic regression to jointly predict composition and its associated uncertainty. To estimate prediction uncertainty, the composition model was formulated as a heteroscedastic Gaussian regression model. Instead of predicting only the mean composition of each component, the network produced two outputs for each component. The first output represented the predicted mean concentration, and the second output represented the predicted uncertainty associated with that concentration. The uncertainty output was constrained to be positive using a positive activation function, so that it could be interpreted as a component-specific standard deviation. This formulation allows the model to learn input-dependent uncertainty, also referred to as heteroscedastic aleatoric uncertainty. This is useful because experimental spectra may contain different levels of residual noise, baseline mismatch, or disagreement with the simulated training distribution even after spectral standardization.

A step-by-step description of setting up an ML model for quantitative prediction from Raman spectroscopy is provided below:

1. Raman spectra of individual components are collected.
2. Equations 1 and 2 are used to generate a simulated spectral library. To improve coverage of the high-dimensional compositional space for multicomponent mixtures, Dirichlet sampling is employed. The resulting dataset is used to train the ML model.
3. Multicomponent mixtures are prepared experimentally, and their Raman spectra are acquired.
4. A correction function is derived by comparing the spectra of experimentally prepared mixtures with those of simulated mixtures of identical composition. Experimental spectra are then corrected using this function relative to the simulated spectra.
5. To validate, test the trained model and correction function on an unseen test dataset.

## Conclusions

ML-assisted Raman spectroscopy is emerging as a powerful analytical tool; however, its broader application is often limited by the availability of high-quality training datasets.^19^ We generated a simulated spectral library by linearly combining pure-component spectra and used it to predict the compositions of highly complex experimentally prepared mixtures. Although the simulated spectra are representative, they do not fully capture the fine variations in experimental data that arise from experimental errors and instrument variation. To address these challenges, we applied a correction strategy that improved the alignment between simulated and experimental spectra, enabling more precise composition prediction in highly complex mixtures.

We demonstrate that the compositions of 15 lipid mixtures were successfully predicted using a machine learning model trained solely on a simulated dataset, achieving R^2^ values exceeding 0.95 for each lipid. This approach minimizes the need for exhaustive calibration datasets while maintaining high quantitative accuracy. The proposed framework offers a practical path for quantitative Raman analysis of chemical mixtures and can easily be extended to quality control applications across various industries.

## Author Information

## Author Contributions

A.Y., A.K., and H.W. contributed to the study design. A.Y. and D.D. performed experiments. A.Y. and A.K. conducted data analysis. A.Y., A.K., and H.W. contributed to data interpretation and drafted the manuscript. All authors reviewed the manuscript.

## Notes

The authors declare no competing financial interests.

## Acknowledgements

Research reported in this publication was supported by the National Institute of General Medical Sciences of the National Institutes of Health under award number (R35GM156609). Table of Contents Graphic was created in BioRender and edited in Adobe Illustrator. Created in BioRender. Yadav, A. (2026). https://BioRender.com/04hb70s. Parts of Figure1 were created in BioRender. Khambhawala, A. (2026). https://BioRender.com/i8wm8jl.

