## Supplementary Information for "Machine Learning-Assisted Raman Spectroscopy for Quantitative Prediction of Multicomponent Mixtures"

**Figure S4. Principal component 1 (PC1) loading vector obtained from PCA of the normalized Raman spectra of the 15 lipids.** PC1 accounts for 45.95% of the total spectral variance. The loading profile highlights Raman regions that contribute most strongly to component separation, with prominent positive contributions near ~1090, ~1260, ~1430, and ~1650 cm⁻¹, and negative contributions near ~930, ~1130, and ~1740 cm⁻¹. Positive and negative loadings indicate spectral features associated with opposite directions along PC1........ 5

**Figure S5 Principal component 2 (PC2) loading vector obtained from PCA of the normalized Raman spectra of the 15 lipids.** PC2 accounts for 14.23% of the total spectral variance. The loading profile identifies spectral regions that provide secondary differentiation among the components, with prominent contributions near ~850–900, ~1050–1100, ~1260, ~1420, ~1660, and ~1740 cm⁻¹. Positive and negative loadings represent spectral features that contribute in opposite directions along PC2. ........ 6

**Figure S6 Cosine similarity matrix comparing the normalized Raman spectra of all pure components.** Each cell represents the cosine similarity between a pair of lipid Raman spectra. Higher values indicate greater similarity in overall spectral shape, with 1 indicating identical spectral profiles. Lower values indicate less similar profiles; a value of 0 indicates no directional similarity, whereas −1 indicates completely opposite profiles. The diagonal elements represent self-similarity and therefore have a score of 1........ 7

**Figure S7 Pearson correlation matrix comparing the normalized Raman spectra of all pure lipids.** Each cell represents the cosine similarity between a pair of lipid Raman spectra. Higher values indicate greater similarity in overall spectral shape, with 1 indicating perfectly positive correlation between spectral patterns. Lower values indicate less similar profiles; a value of 0 indicates no linear correlation between the spectral intensity variations, whereas −1 indicates perfectly negatively correlated spectral patterns. The diagonal elements represent self-similarity and therefore have a score of 1. ........................................................................................ 8
**Figure S8 Comparison between normalized Raman spectra of GTL and OlA.**...................... 9
**Figure S9 Comparison between experimental Raman spectra and simulated spectra.** (a) Kernel-density contours of PCA score distributions for the experimental and simulated datasets, showing mismatches in the PC1-PC2 space. (b) Normalized experimental and simulated Raman spectra, showing differences in the overall spectral profile. (c) Comparison of PCA score profiles across principal components for experimental and simulated data, illustrating their mismatches. ......................................................................................................................................... 10
**Figure S10 Visualization of spectral mismatch.** Comparison between normalized experimental and simulated spectra, with highlighted regions showing major intensity differences. .............. 11

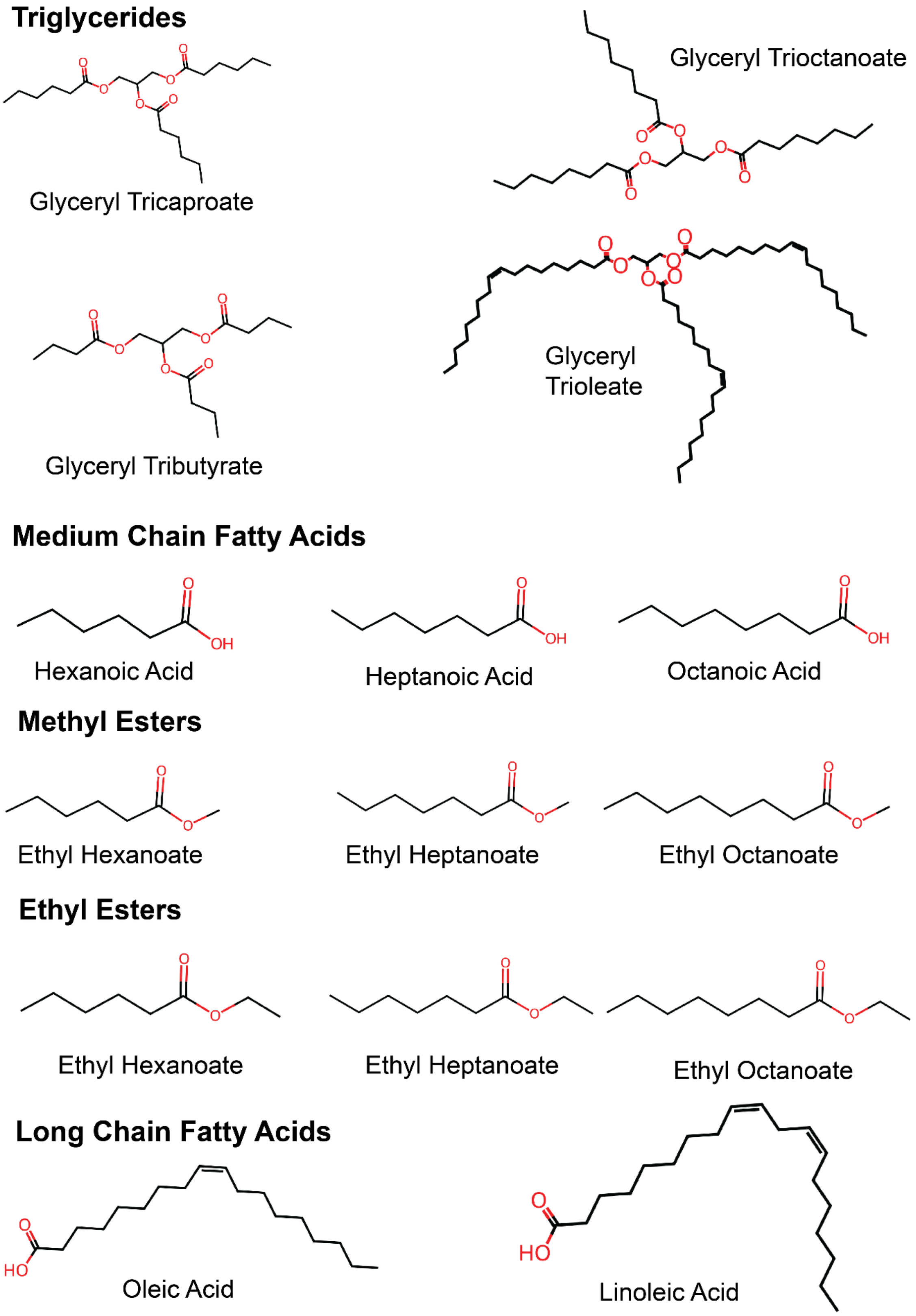

**Figure S1 Molecular structures of lipids used in the study.** The selected lipids include 3 groups; triglycerides, fatty acids and esters.

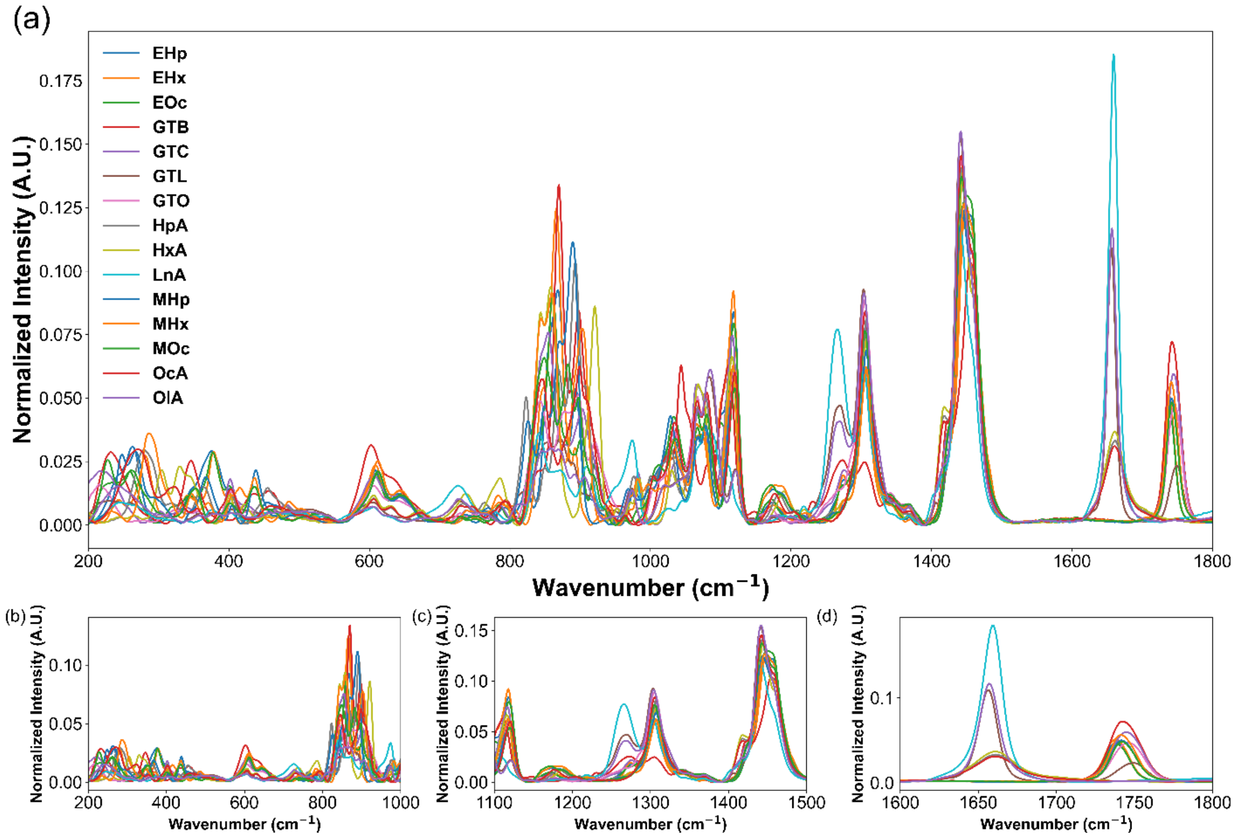

**Figure S2 Normalized Raman spectra of the 15 lipids.** (a) Overlay of the pure-component Raman spectra, spectral range of 200–1800 cm$^{-1}$. (b-d) Expanded views of selected spectral regions: (b) 200–1000 cm$^{-1}$; (c) 1100–1500 cm$^{-1}$; and (d) 1600–1800 cm$^{-1}$.

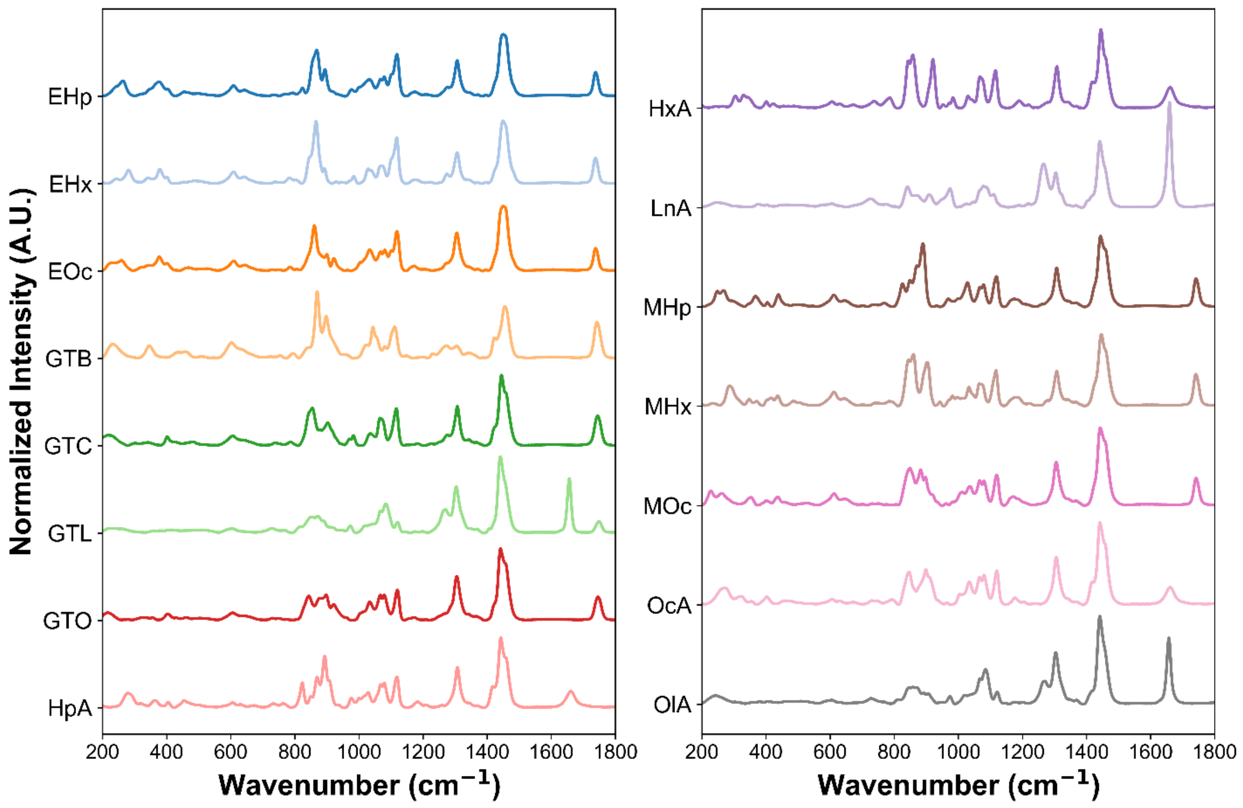

**Figure S3 Stacked, normalized Raman spectra of the lipids used in this study.**

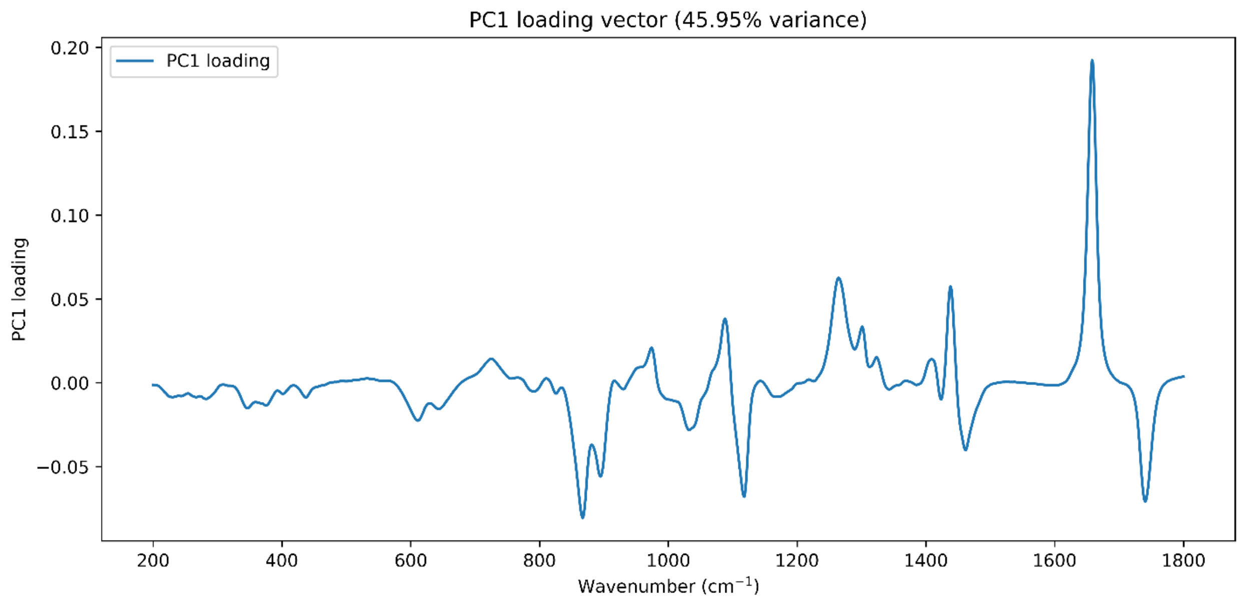

**Figure S4. Principal component 1 (PC1) loading vector obtained from PCA of the normalized Raman spectra of the 15 lipids.** PC1 accounts for 45.95% of the total spectral variance. The loading profile highlights Raman regions that contribute most strongly to component separation, with prominent positive contributions near ~1090, ~1260, ~1430, and ~1650 cm⁻¹, and negative contributions near ~930, ~1130, and ~1740 cm⁻¹. Positive and negative loadings indicate spectral features associated with opposite directions along PC1.

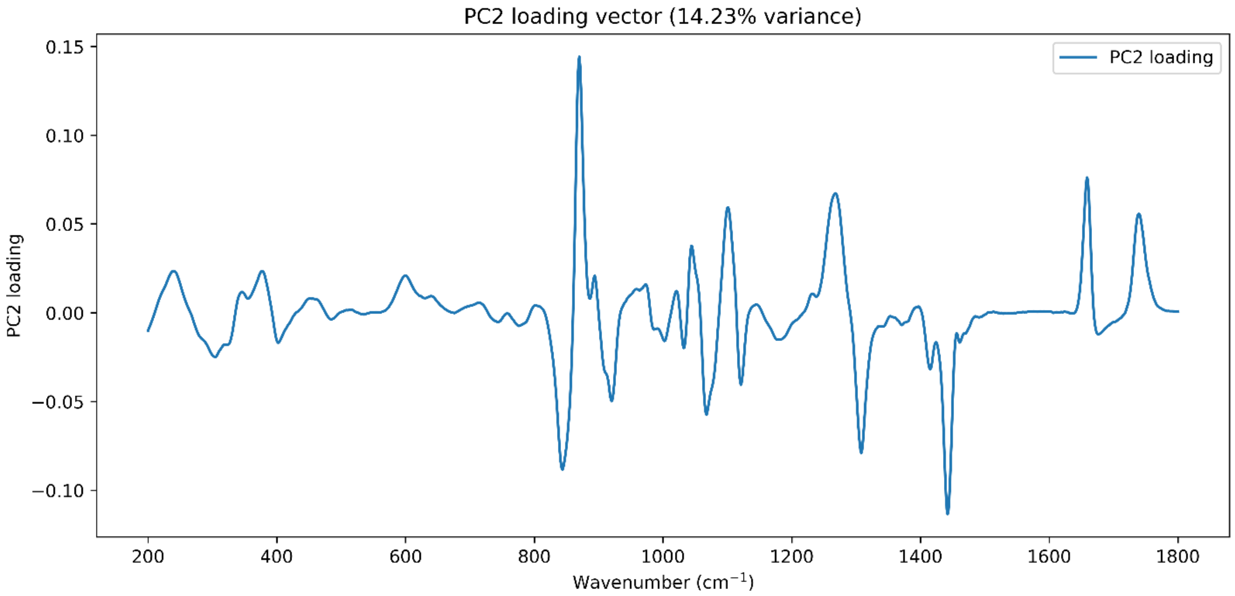

**Figure S5 Principal component 2 (PC2) loading vector obtained from PCA of the normalized Raman spectra of the 15 lipids.** PC2 accounts for 14.23% of the total spectral variance. The loading profile identifies spectral regions that provide secondary differentiation among the components, with prominent contributions near ~850–900, ~1050–1100, ~1260, ~1420, ~1660, and ~1740 cm⁻¹. Positive and negative loadings represent spectral features that contribute in opposite directions along PC2.

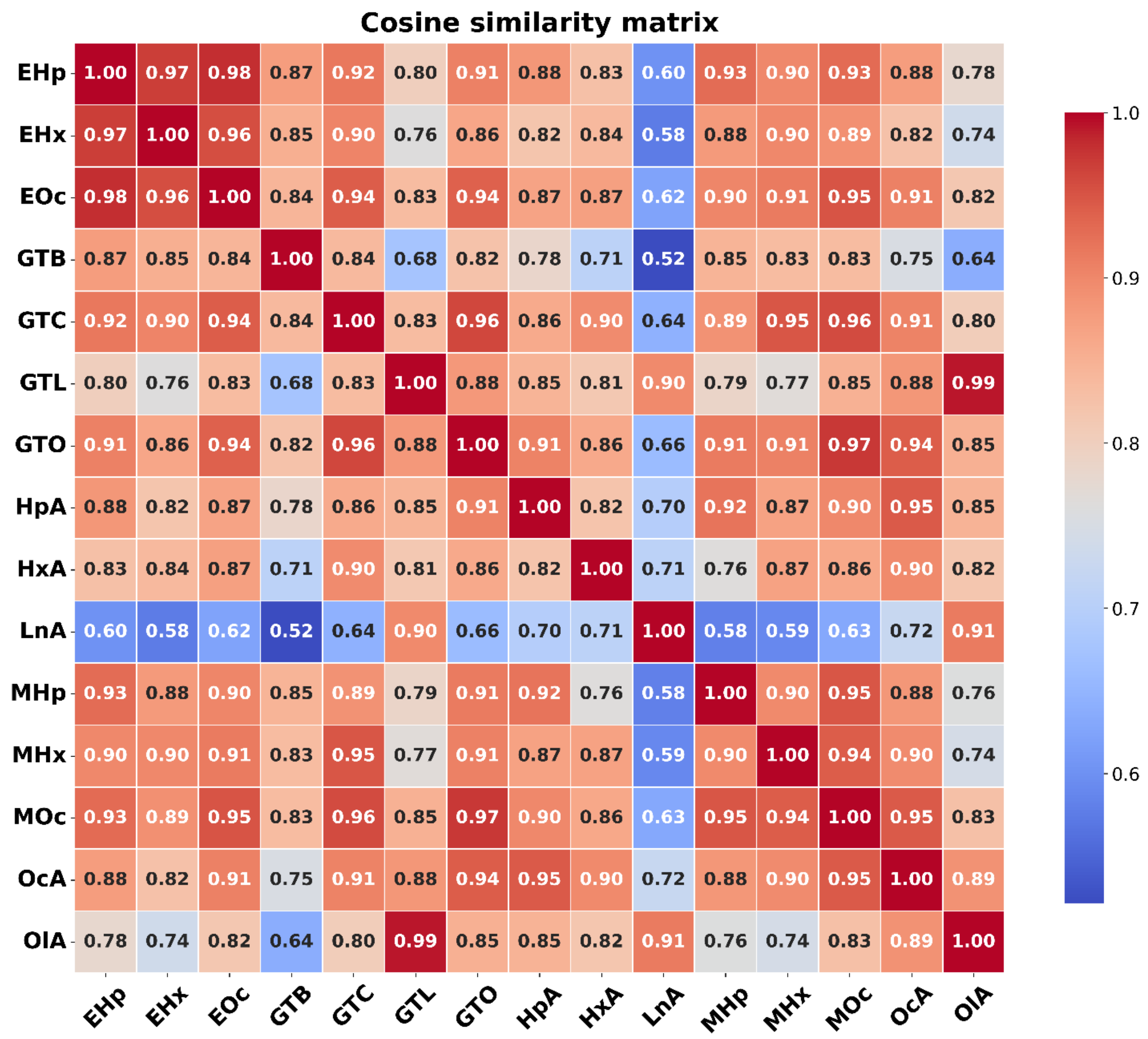

**Figure S6 Cosine similarity matrix comparing the normalized Raman spectra of all pure components.** Each cell represents the cosine similarity between a pair of lipid Raman spectra. Higher values indicate greater similarity in overall spectral shape, with 1 indicating identical spectral profiles. Lower values indicate less similar profiles; a value of 0 indicates no directional similarity, whereas −1 indicates completely opposite profiles. The diagonal elements represent self-similarity and therefore have a score of 1.

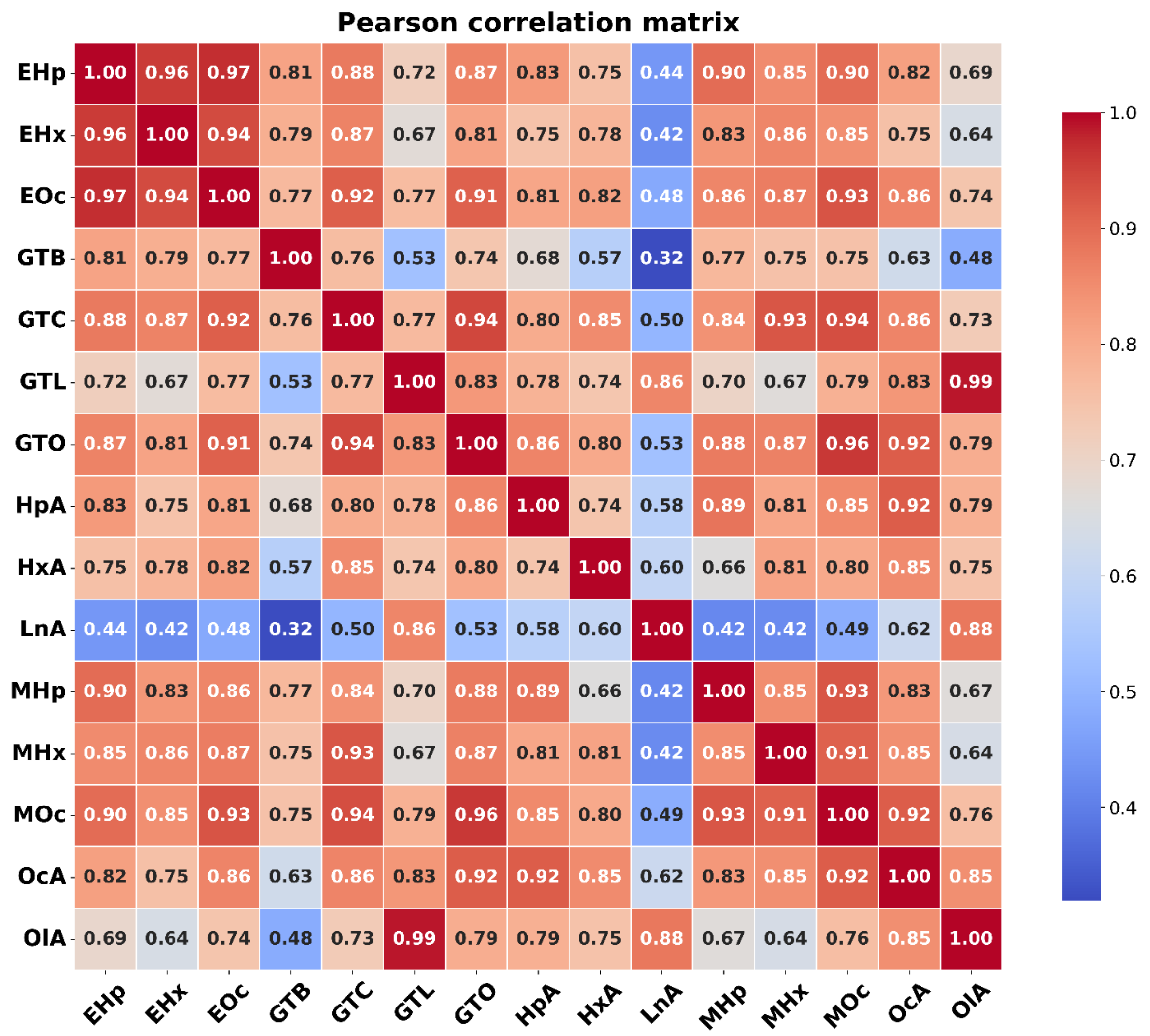

**Figure S7 Pearson correlation matrix comparing the normalized Raman spectra of all pure lipids.** Each cell represents the cosine similarity between a pair of lipid Raman spectra. Higher values indicate greater similarity in overall spectral shape, with 1 indicating perfectly positive correlation between spectral patterns. Lower values indicate less similar profiles; a value of 0 indicates no linear correlation between the spectral intensity variations, whereas −1 indicates perfectly negatively correlated spectral patterns. The diagonal elements represent self-similarity and therefore have a score of 1.

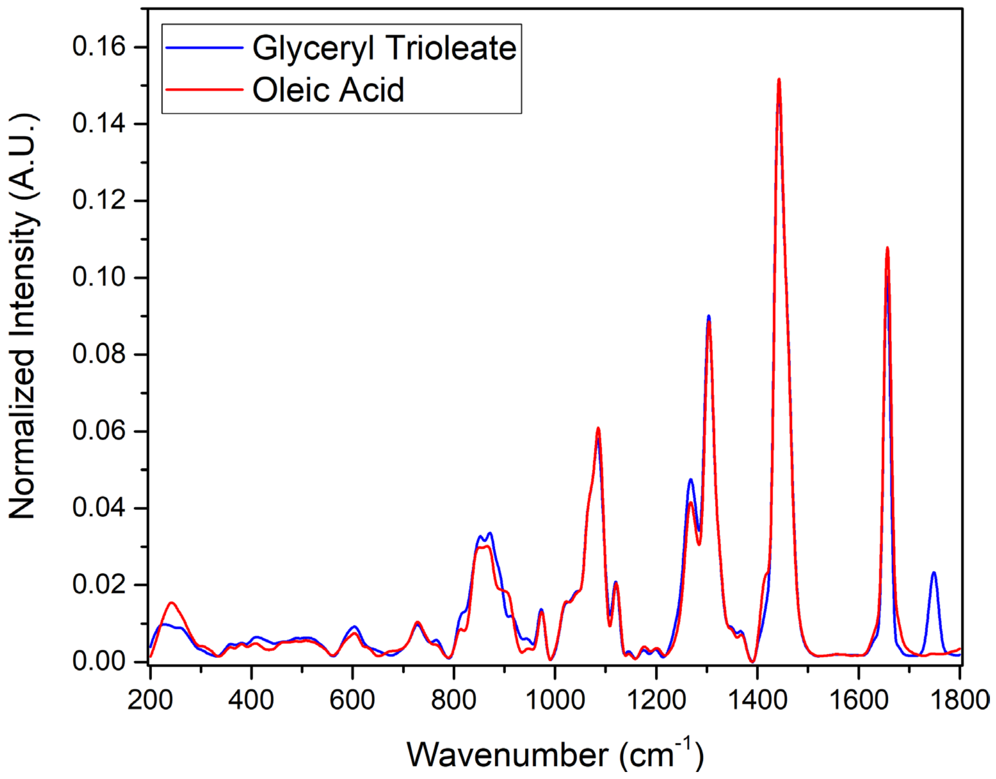

**Figure S8 Comparison between normalized Raman spectra of GTL and OlA.**

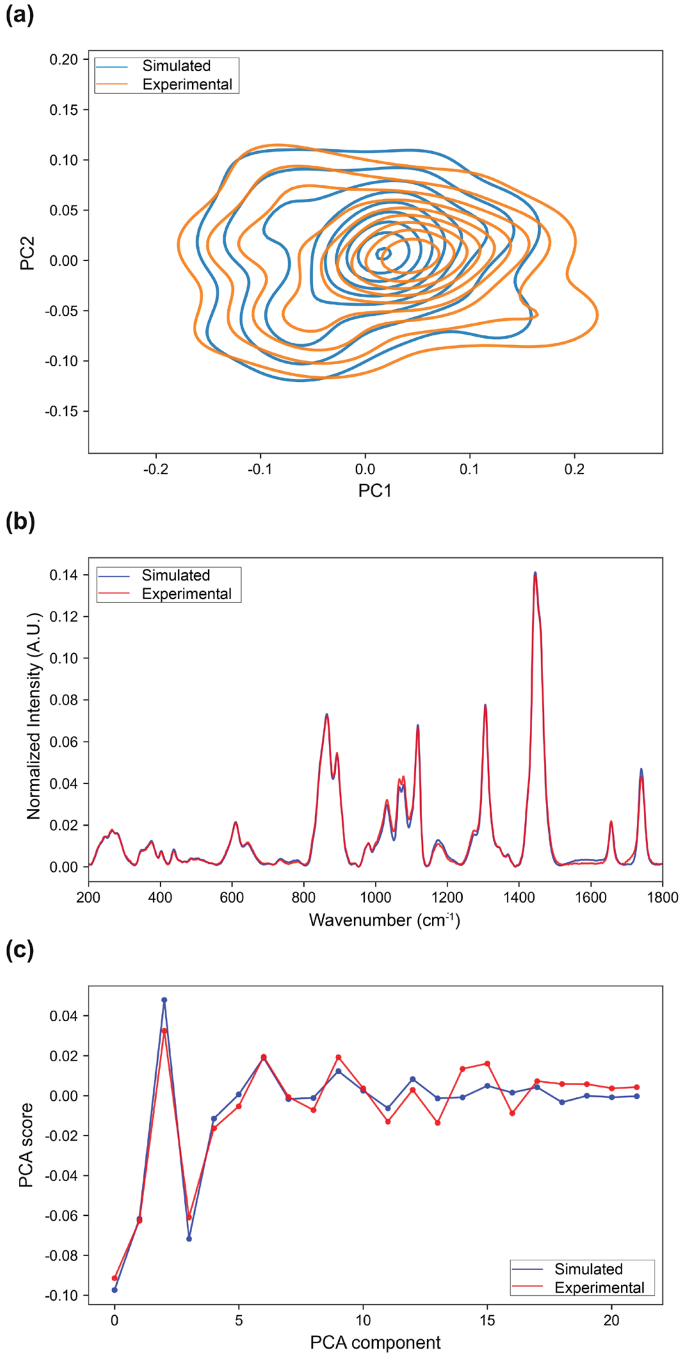

**Figure S9 Comparison between experimental Raman spectra and simulated spectra.** (a) Kernel-density contours of PCA score distributions for the experimental and simulated datasets, showing mismatches in the PC1-PC2 space. (b) Normalized experimental and simulated Raman spectra, showing differences in the overall spectral profile. (c) Comparison of PCA score profiles across principal components for experimental and simulated data, illustrating their mismatches.

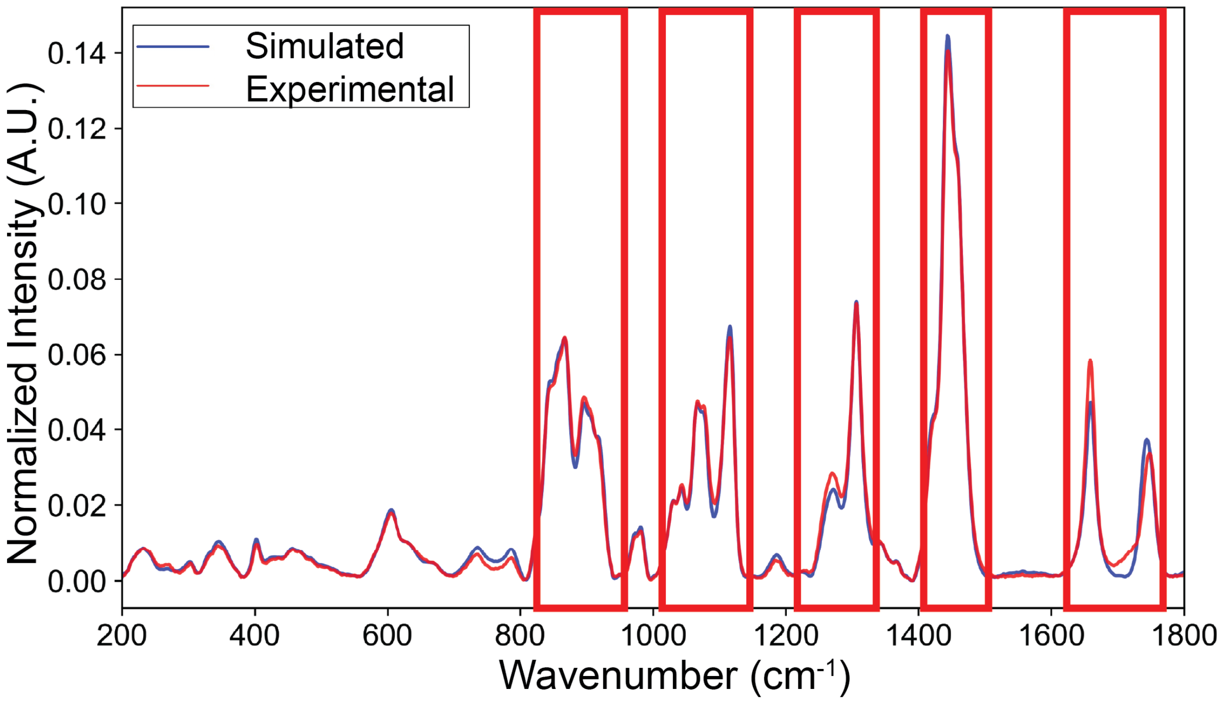

**Figure S10 Visualization of spectral mismatch.** Comparison between normalized experimental and simulated spectra, with highlighted regions showing major intensity differences.
